# Modifying and reacting to the environmental pH drives bacterial interactions

**DOI:** 10.1101/136838

**Authors:** Christoph Ratzke, Jeff Gore

## Abstract

Microbes usually exist in communities consisting of myriad different but interacting species. These interactions are typically mediated through environmental modifications; microbes change the environment by taking up resources and excreting metabolites, which affects the growth of both themselves and also other microbes. A very common environmental modification is a change of the environmental pH. Here we show that changing and reacting to the pH leads to a feedback loop that can determine the fate of bacterial populations. We find experimentally that these pH changes can decide the fate of bacterial populations; they can either facilitate or inhibit growth, and in extreme cases will cause extinction of the bacterial population. Moreover, modifying the pH can determine the interactions between different species. Understanding how single species change the pH and react to it allowed us to estimate their pairwise interaction outcomes. Those interactions lead to a set of generic interaction motifs—bistability, succession, murder suicide, and mutualism—that may be independent of which environmental parameter is modified and thus reoccur in different microbial systems.

## Introduction

Microbes thrive essentially everywhere on this planet, usually as part of complex multi-species communities^1^. The interactions between the microbes influence their growth and survival and thus the composition of those communities^2–6^. Although microbial interactions are of clear importance, only limited insights exist. They are often obtained from microbial systems where a specific molecular mechanism causes a specific interaction, like mutualism caused by cross-feeding of amino acids^7–11^, cross-protection from antibiotics ^12^ or competition by toxins ^13,14^.

Despite the large number of possible types of interactions there are certain points they all have in common, as interactions are in general mediated through the environment. First, microbes change the environment by consuming resources and excreting metabolites. Second, these changes to the environment influence the growth and survival of both the microbe that originally altered the environment as well as other microbial species that are present. Thus the interaction between microbes may be set by how their metabolisms change the environment and react to it. A very important parameter for microbes is the pH, and different species prefer different pH values. Thus pH strongly influences the species composition in soil^15–17^ or the human gut microbiome^18^. On the other hand, many biochemical reactions involve a turnover of protons and thus microbes also alter the pH around them. We show here that the way microbes modify the pH of their environment feeds back on them but also influences other microbes. This determines their growth behavior and the interactions between different bacterial species based upon how each species interacts with pH changes in the environment.

## Results

Growing a collection of soil microbes in a medium that contains glucose as main carbon source and urea as main nitrogen source leads to a change of the initial pH for nearly all tested bacteria (n=119) (Fig. 1a). Therefore microbial growth modulates the pH, but the pH is also known to strongly affect microbial growth. In this way microbes feed back on their own growth, but also influence the growth conditions of other species that might be present (Fig. 1b).

**Figure 1:**
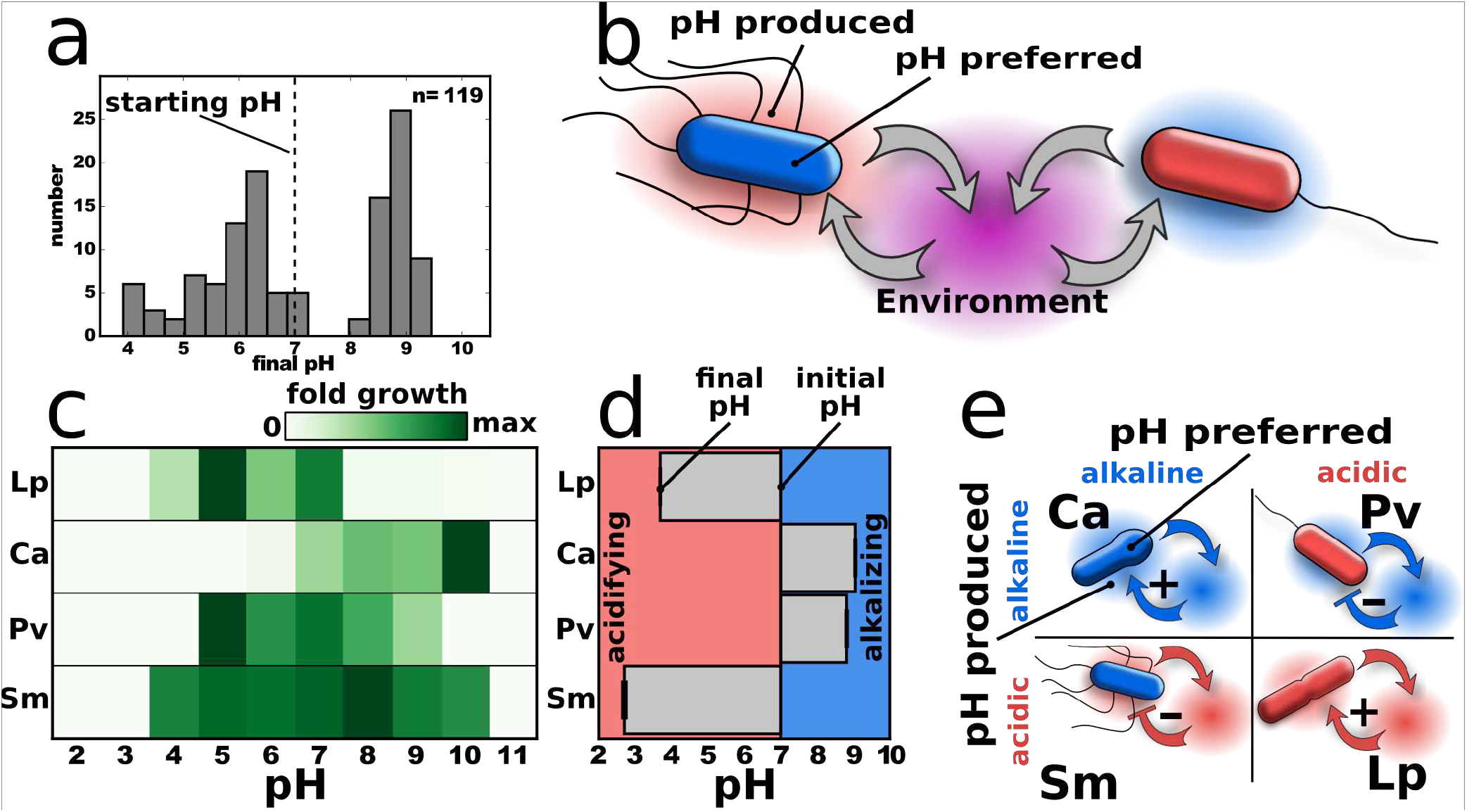
*Microbes modify the environment and react to it. (a)* *A collection of soil bacteria grown in a medium that contains urea and glucose can lower or increase the pH starting at pH 7 (dashed line). Also growing the soil bacteria in Luria-Bertani medium causes pH changes (Supplementary Fig. 1 e)* ***(b)*** *By changing the environment bacteria influence themselves but also other microbes in the community.* ***(c)*** Lp and Pv *prefer acidic, Ca prefers alkaline and Sm has a slight preference towards alkaline environments. Shown is fold growth in 24h. The bacteria are grown on buffered medium with little nutrient to minimize pH change during the growth (Methods and Supplementary Fig. 1c).* ***(d)*** *Starting at pH 7 Lp and Sm decrease and Ca and Pv increase the pH (Supplementary Fig. 3). Only little buffering, 10g/L glucose and 8g/L urea as substrates were used in* ***(d). (e)*** *Microbes can increase or decrease the pH (blue environment is alkaline and red environment is acidic) and thus produce a more or less suitable environment for themselves. Blue bacteria prefer/tolerate alkaline and red acidic conditions.*

Microbes can lower or increase the pH, which may be beneficial or deleterious for their own growth. This leads to four possible combinations, for which we identified an example species of each (Fig. 1b,c,d; Methods and Supplementary Fig. 1 and 2). *Lactobacillus plantarum* (Lp) is an anaerobic bacterium that produces lactic acid as metabolic product and thus lowers the pH, but also prefers low pH values^19^. *Corynebacterium ammoniagenes* (Ca) produces the enzyme urease that cleaves urea into ammonia and thus increases the pH^20^; at the same time it prefers higher pH values. *Pseudomonas veronii* (Pv) also increases the pH of the medium, but prefers low pH values for growth. Finally, *Serratia marcescens* (Sm) strongly lowers the pH^21^ but better tolerates comparably higher pH values, with a slight optimum at around pH 8. As expected, the strength of the pH change depends on the amount of glucose and urea (Supplementary Fig. 1a) and can be tempered by adding buffer (Supplementary Fig. 1b). A pH change can also be first good and then bad for a microbe (Supplementary Fig. 1d), however we focus here on the simple cases. In summary, we find that microbial growth often leads to dramatic changes in the pH of the environment, and this pH change can promote or inhibit bacterial growth.

When the pH modification is beneficial for the bacteria there is a positive feedback on their growth. The more bacteria there are, the stronger they can change the environment and thus the better they do. At adverse pH conditions a sufficiently high cell density may therefore be needed to survive at all – an effect known as strong Allee effect^22,23^. Indeed we observe such an effect in our bacteria: Ca promotes its own growth by alkalizing the environment, leading to a minimal starting cell density required for survival under daily batch culture with dilution (Fig. 2a). Adding buffer (Fig. 2a, right) or lowering the nutrient concentration (Supplementary Fig. 4) tempers the pH change and thus necessitates an even higher bacterial density for survival. Changes to the pH could therefore be a common mechanism of cooperative growth that leads to an Allee effect and an associated minimal viable population size.

**Figure 2:**
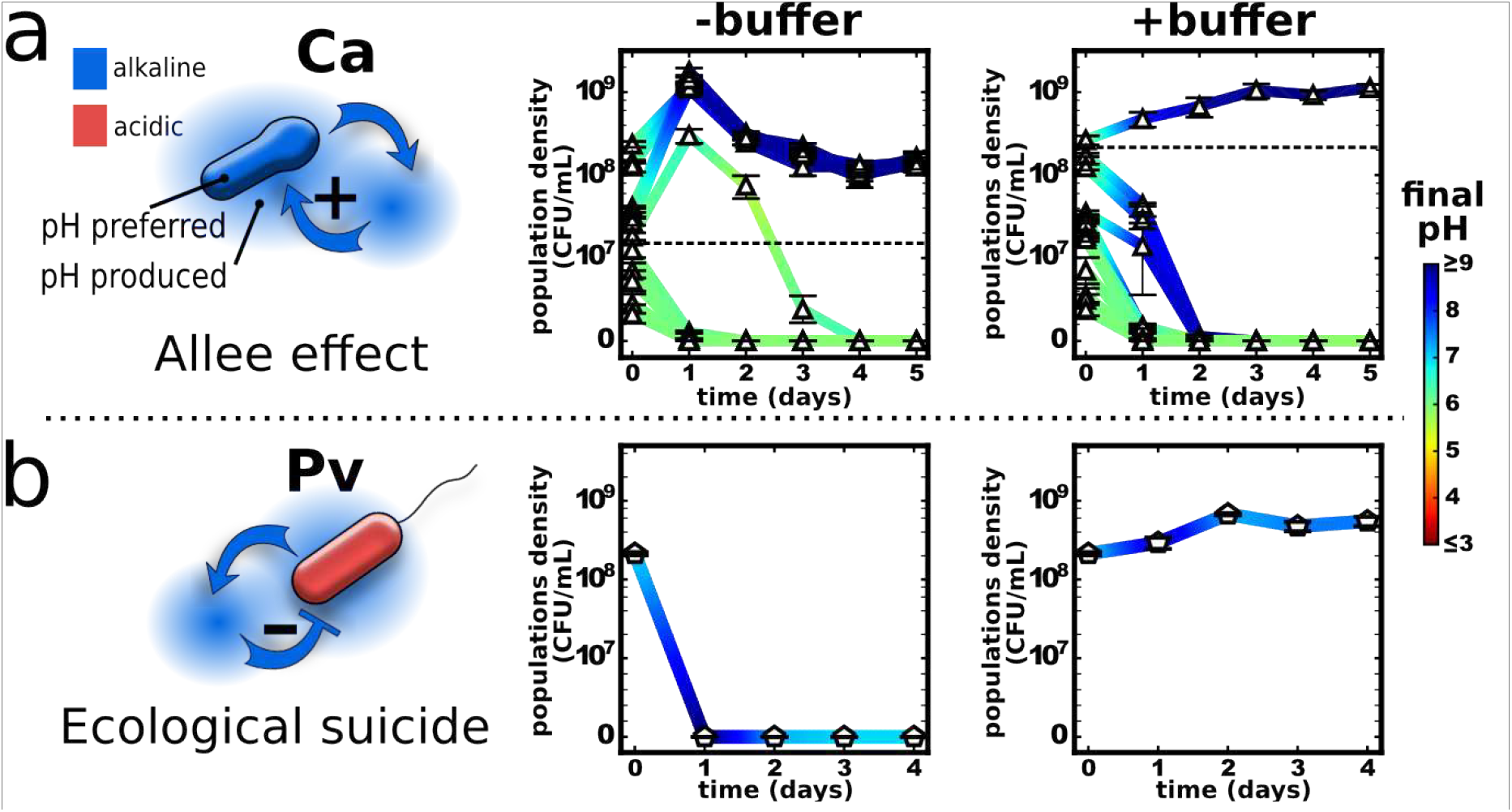
Single species can enhance or inhibit their own growth via changing the pH. The curves show bacterial density over time and the color shows the pH. **(a)** Ca increases the pH and also prefers these higher pH values, leading to a minimal viable cell density required for survival. Increasing the buffer concentration from 10mM to 100mM phosphate makes it more difficult for Ca to alkalize the environment and therefore increases the minimal viable cell density. **(b)** Pv also increases the pH, yet prefers low pH values. Indeed, Pv populations can change the environment so drastically that it causes the population to go extinct. Adding buffer tempers the pH change and thus allows for the survival of Pv. An Allee effect can also be found in Lp and ecological suicide in Sm (Supplementary Fig. 6).

Bacteria can also change the environment in a way that harms themselves, for example by shifting the pH away from the growth optimum. In the most extreme case the bacteria may make the environment so detrimental that it becomes deadly for them, an effect we call ecological suicide. Indeed, we find that Pv, which prefers lower pH, alkalizes the medium and thus causes its own extinction (Fig. 2b). Again adding buffer (Fig. 2b, right) or lowering nutrient concentrations (Supplementary Fig. 5a) tempers the pH change and saves the population. It is therefore not the initial condition that kills the bacteria, but the way the population changes the pH, which is further underlined by OD measurements and following the CFU over time (Supplementary Fig. 5b and c). The surprising effect of ecological suicide is investigated in more detail in a separate manuscript.

Microbes that modify the environment not only affect their own growth but also other microbial species that may be present. In this way the environmental modifications could drive interspecies interactions. We described previously (Fig 1) that there are four different ways that a single species can change the environment and in turn be affected: lowering and increasing the pH, and affecting the growth in positive or negative ways. Accordingly, there exist in principle 6 (4 choose 2) pairwise combinations of the four types in Fig. 1d; however, two of them are symmetric cases of each other, leaving 4 unique interaction types (Supplementary Fig. 7). We have seen that the microbial metabolism can determine the fate of a population by changing and reacting to the pH. Thus can we in a similar way understand and perhaps even anticipate possible outcomes of those four interactions based on the properties of the single species? To explore this question we captured the essential elements of the pH interaction via a simple differential equation model describing the growth of the two species (see Supplementary Information and Supplementary Fig. 8):

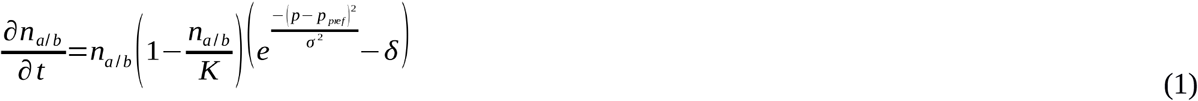

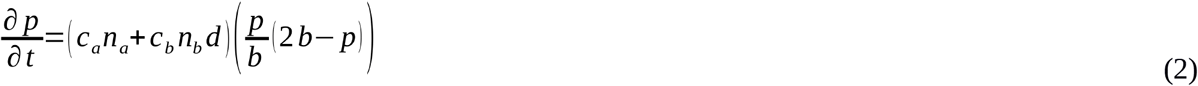

The bacteria densities n_a/b_ follow a logistic growth that saturates at the carrying capacity *K*, but growth becomes maximal at a preferred proton concentration p_pref_. The further away the proton concentration is from the optimum of the species, the slower the bacteria grow and finally start to die. The proton concentration p is changed by the bacteria, according to their density n_a/b_ and their ability to change the proton concentration c_a/b_ which is set to +/- 0.1. Multiplication by a quadratic function ensures that the proton concentration stays within [0,2b].

For the single species cases of Lp and Ca this model naturally gives an Allee effect – a minimal initial bacterial density is needed for survival depending on the initial pH value (Fig. 3 and Supplementary Fig. 8 f, g). In addition, when the species are changing the environment in a way that is bad for them (eg Pv and Sm), the model yields ecological suicide in which the population dies under all starting conditions (Fig. 3 and Supplementary Fig. 8 h,i). Thus, this simple model can capture the measured outcomes for the single species.

**Figure 3:**
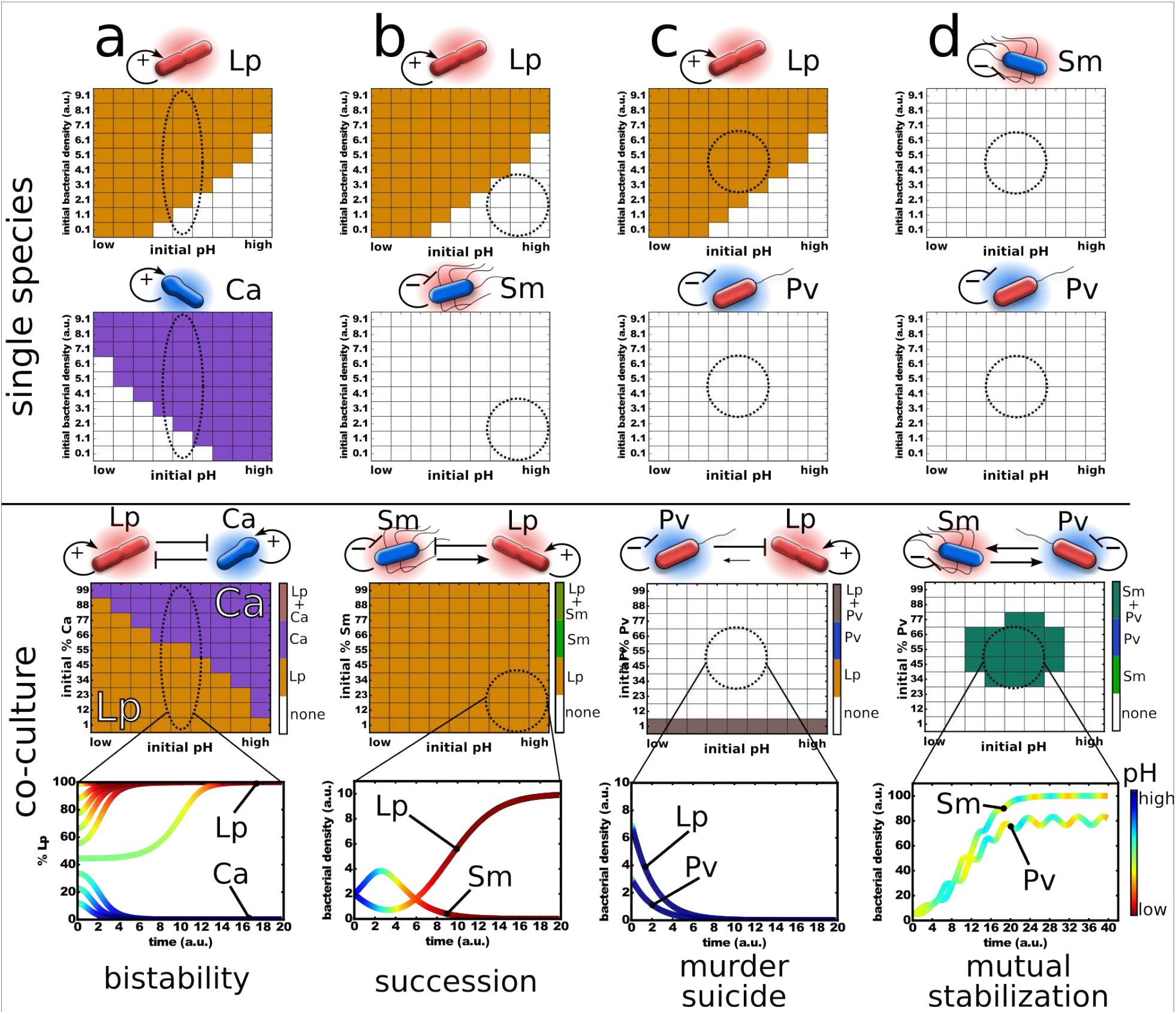
The metabolic properties of the different species may allow to estimate their interactions. A model based on differential equations was set up to simulate the bacterial growth (see main text). The survival of the species for different initial parameter values is plotted in the following. The upper two panels for each column show the simulation outcome for the single species. The panel below the outcomes for the co-cultures and the lowest panel the dynamics over time in the region of the phase diagrams marked by the dashed circles. The “pH” scale reaches from low to high which corresponds to a “proton concentration” of 10 to 0. s was set to 4 and d to 0.5. c_a/b_ was set to +/- 0.1, b to 5 and d as described in the supplement. (a) Lp and Ca show bi-stability in co-culture depending on the initial fraction and pH. (b) Lp and Sm show succession at high initial pH values, where Lp can only survive if the pH was first lowered by Sm. (c) If Pv lowers the proton concentration strong enough it can kill itself and also Lp, resulting in murder suicide. (d) Sm and Pv can protect each other from ecological suicide and coexist, whereas they can not survive on their own. A fuzzy logic based model leads to very similar results (Supplementary Information and Supplementary Fig. 9 and 10).

We next explored whether the model can provide insight into the two species interactions. In particular, there is a question of whether just one environmental parameter - the pH - is sufficient to predict non-trivial outcomes of interspecies competition, neglecting all the other ways in which the species interact. In the following we focus on the especially interesting outcomes predicted by the model (marked by dashed circles in Fig. 3). To test our predictions, those four different interaction pairs were grown with daily dilutions both as single species and in pairs under the conditions suggested from the model (Fig. 4).

**Figure 4:**
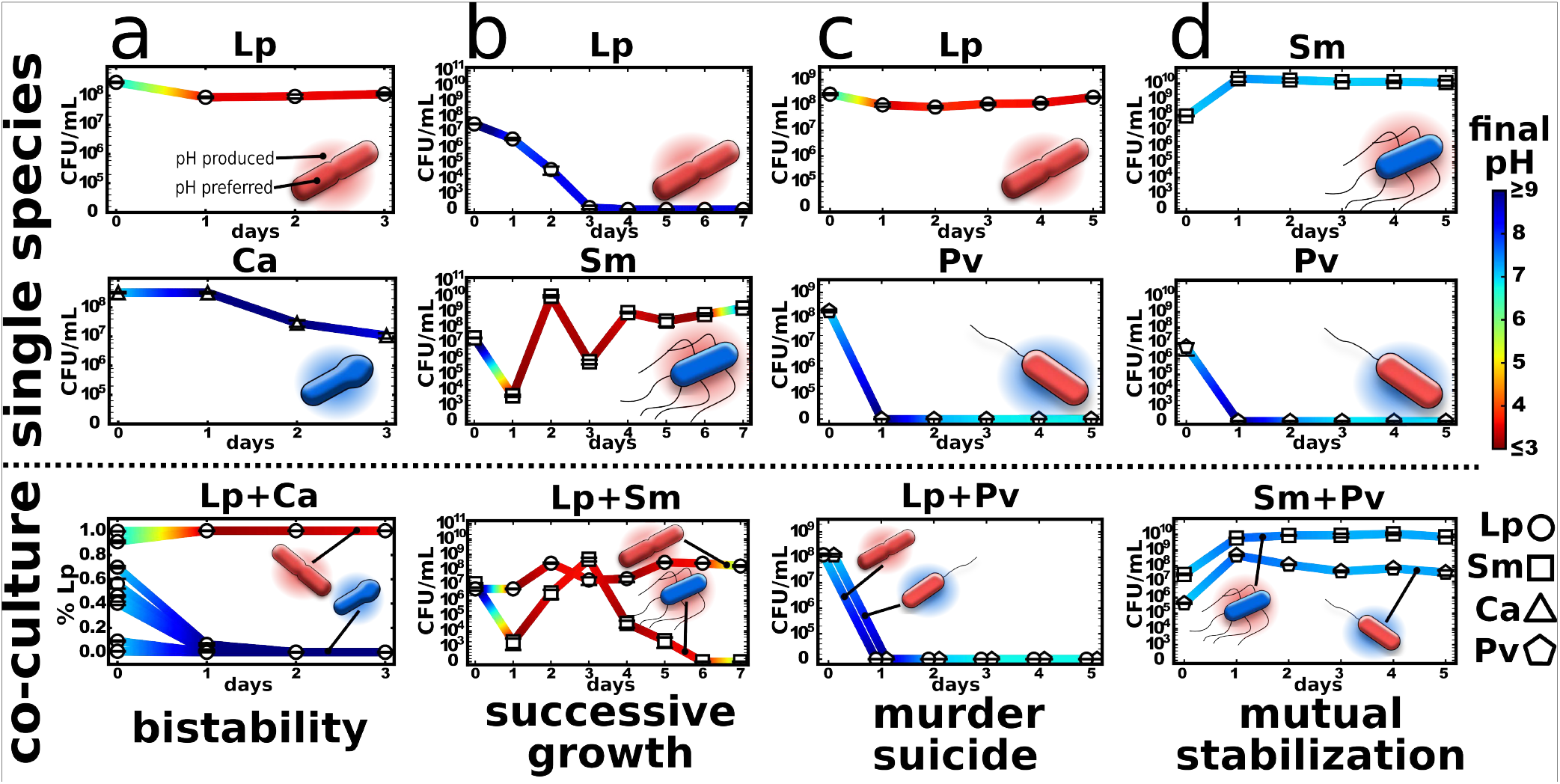
Changing the environment drives interactions between microbes. Four different interaction types can be found depending on how the environmental changes act on the organisms themselves and each other. **(a)** Lp and Ca produce bi-stability. **(b)** Sm and Lp show successive growth. **(c)** Pv commits murder suicide on Lp. **(d)** Sm can stabilize Pv, when the medium is sufficiently buffered. For a more detailed description of the different interactions cases see text. The media composition and protocols are slightly different for the different cases. See Methods and Supplement for details.

Lp and Ca each modify the environment in a way that is good for themselves yet bad for the other species. Given this “antagonistic niche construction”, the model predicts that the co-culture can yield bistability, in which only one species will survive, whereas the winner depends on the initial species abundances and initial pH (Fig. 3a). Consistent with this prediction, we experimentally find bistability in the co-culture of Lp and Ca, with the initially abundant species driving the other species extinct (Fig. 4a and Supplementary Fig. 11 b-d). As expected, removing glucose and urea from the medium prevents pH changes and thus also removes the forces leading to bi-stability, which results in coexistence between the two species (Supplementary Fig. 11a). Two species that are attempting to modify the environment in incompatible ways can therefore lead to the emergent phenomenon of bistability.

Lp can not tolerate high pH values, whereas Sm can initially grow at high pH, but acidifies the environment and kills itself (Supplementary Fig. 6 and Supplementary Fig. 8 g,i). This raises the question of whether Sm can aid Lp in surviving high pH values. Indeed, our model suggests that it may be possible to observe succession, in which Sm grows first and lowers the pH, thus allowing Lp to survive in a condition that it would not have been able to survive on its own (Fig. 3b and Supplementary Fig. 8g,i). We can indeed find this interaction motif experimentally. At an initially high pH Lp can only survive if Sm helps to modify the environment (Fig. 4b). However, contrary to the simulation Sm alone does not experience ecological suicide in the experiment, likely because the high starting pH prevents Sm from lowering the pH too rapidly. Indeed, at lower starting pH values Sm shows ecological suicide (Supplementary Fig. 6 and Supplementary Fig. 8e). Adding buffer or removing nutrient causes only moderate acidification and thus kills Lp but allows Sm to survive (Supplementary Fig. 12). The need to alter the pH of the media can lead to something akin to altruism between species, where the “selfless” species does not survive the ecological interaction.

In the previous paragraph we considered the case of a species transiently helping another species to survive, raising the question of whether it might also be possible to observe the opposite situation in which pH modification leads to ecological suicide and kills the interactions partner with it. Indeed, our model predicts that this “murder-suicide” could be present for the Lp/Pv pair over a wide range of cell densities and starting pH values (Fig 3c). To explore this prediction of our model experimentally, we utilize Pv, which commits ecological suicide via alkalization (Fig 2b), together with Lp, which dies in alkaline conditions (Fig 2a). Once again consistent with the predictions of our simple model, we successfully observed such a murder-suicide, in which the extreme alkalization of the media by Pv led to the death of both Pv and Lp (Fig 4c). The addition of buffer tempers this alkalization sufficiently to stop Pv from suicide but still allows it to out-compete Lp (Supplementary Fig. 13). However, if Lp were more effective at acidifying the media then coexistence might have been possible (Supplementary Fig. 8a). We therefore find that ecological suicide can also have a negative impact on other present species.

Finally, the model suggests that it may be possible for two species to help each other avoid ecological suicide (Fig. 3d). This would correspond to an obligatory mutualism, in which the opposing the pH changes cancel each other and therefore allow for the stable coexistence of the two species in an environment in which neither species could survive on its own. However this mutual stabilization is only observed in a rather small parameter range, which suggest that finding it experimentally may be difficult. Indeed, experimentally we could not find conditions in which Pv and Sm form the most extreme form of mutual stabilization, in which each species alone results in ecological suicide yet together they survive. However, we did observe a situation in which Pv alone causes ecological suicide yet Sm can stabilize the pH and allow both species to survive and coexist (Fig. 4d; see methods for details). Species that harm themselves through their environmental modifications can therefore benefit from other species in the environment.

## Discussion

Although bacterial interactions are commonly regarded as a very complex business, we found here a surprising simplicity. By studying how bacteria change their environment and react to it we could understand a variety of interactions within populations as well as interactions between different species. Although the interaction outcomes were quite different they were all mediated by a single environmental parameter - the pH - which shows that even a rather complex set of interactions can be mediated through the same environmental parameter. This significantly simplifies the situation and gives hope that even more complex microbial communities may be tractable. Moreover the pH is a very general parameter that essentially all microbes influence and depend on, which makes it likely that interactions through pH modification happen rather often in nature.

Many microbes are known to show an Allee effect by cooperatively secreting ‘public goods’ - like enzymes that break down complex sugars^24–27^. However, in our system the Allee effect was mediated by the pH without the necessity of a specialized enzymatic machinery. This suggests that cooperation may be easier to achieve and more widespread than is often assumed.

Cooperation is a very important collective phenomenon, where organisms work together to achieve something they could not on their own. However, we could also find a different type of collective action where bacteria collectively deteriorate their environment and thus cause their own extinction – the ecological suicide. We discuss this surprising effect in more depth in a separate manuscript (in preparation).

After understanding the interactions within a population as mediated through the environment, we went a step further and applied the same reasoning to interspecies interactions. Surprisingly a simple model could quite accurately forecast the interaction motifs that we later found experimentally: bi-stability, successive growth, murder suicide and mutual stabilization - all driven by pH modifications. In our model (Fig. 3) the pH could be replaced by any other environmental parameter that is affected by and affects microbes – like oxygen or metabolite concentrations. Thus describing microbial interactions as a combination of modifying the environment and reacting to it may give a general framework for microbial interactions independent of the exact underlying biochemical mechanism. And indeed several examples for the described interaction motifs have been found in other studies. Bi-stability has been observed in the gut microbiome^28^, successive growth in the colonization of chitin particles^29^ or the human gut^30^ and mutual stabilization in a variety of cross-feeding and cross-protection metabolisms^9,12^. To our knowledge ecological suicide and thus also murder suicide are not yet described in microbes. However, a self-inflicted decline of populations could be found in several macro-organisms^31–33^.

In the age where many genomes are sequenced and metabolic models of microbes can be generated^30,34–37^ measuring metabolic properties of microbes may eventually become obsolete and interactions could be directly predicted based on genome sequences. Together with such approaches our framework for microbial interactions may become an important part of the future goal to predict the assembly, composition and function of complex microbial communities.

## Methods

All chemicals were purchased from Sigma Aldrich, St. Louis, USA, if not stated otherwise.

### Buffer

For precultures of the bacteria the basic buffer recipe was 10g/L yeast extract (Becton Dickinson, Franklin Lakes, USA) and 10g/L soytone (Becton Dickinson, Franklin Lakes, USA). We refer to that buffer as 1xNutrient medium. For the different bacteria the initial pH was either 6 or 7 and in some cases 100mM Phosphate were added, as outlined in the single experiments below. For the washing steps and the experiments itself the medium contained 1g/L yeast extract and 1g/L soytone, 0.1mM CaCl_2_, 2mM MgCl_2_, 4mg/L NiSO_4_, 50mg/L MnCl_2_ and 1x Trace Element Mix (Teknova, Hollister, USA). We refer to that buffer as base buffer. It was supplemented with phosphate buffer and/or glucose and urea as outlined in the single experiments. The glucose and urea was added freshly every day directly before starting the experiments, to avoid degradation of the urea. The usual concentrations were 10g/L glucose and 8g/L urea, deviations from that are described for the single experiments below. All media were sterile filtered.

### Estimation of Colony Forming Units (CFU)

To estimate the number of living bacteria in the different experiments we used colony counting. At the end of every growth cycle a dilution row of the bacteria by diluting them 7 times 1/10x in phosphate buffered saline (PBS, Corning, New York, USA) was made. With a 96-well pipettor (Viaflo 96, Integra Biosciences, Hudson, USA) 10µL of every well for every dilution step were transferred to an agar plate (Tryptic Soy Broth (Teknova, Hollister, USA), 2.5% Agar (Becton Dickinson, Franklin Lakes, USA) and 50mg/L MnCl_2_ in case of Lp was plated) with 150mm diameter. The droplets were allowed to dry in and the plates were incubated at 30°C for 1-2 days until clear colonies were visible. The different dilution steps made sure that a dilution could be found that allows to count single separated colonies. The different bacteria could be distinguished by their colony morphology (Supplementary Fig. 2).

### pH measurements

To measure the pH directly in the bacterial growth culture at the end of each growth cycle a pH microelectrode (N6000BNC, SI Analytics, Weilheim, Germany) was used. The grown up bacterial cultures were transferred into 96-well PCR plates (VWR, Radnor, USA) that allowed to measure pH values in less than 200µL.

### Bacterial culture

All cultures were incubated at 30°C. The precultures were done in 5mL medium in 50mL culture tubes (Falcon/Becton Dickinson, Franklin Lakes, USA) over night in different variations of the 1xNutrient described above. The shaking speed was 250rpm on a New Brunswick Innova 2100 shaker (Eppendorf, Hauppauge, USA), the lids of the falcons tubes were only slightly screwed on to allow gas exchange. The experiments were all done in 96-deepwell plates (Eppendorf, Hauppauge, USA) covered with two sterile AearaSeal adhesive sealing films (Excell Scientific, Victorville, USA), the plates were shaken at 1350rpm on Heidolph platform shakers (Titramax 100, Heidolph North America, Elk Grove Village, USA). To avoid evaporation the shakers were covered with a custom made polyacryl box (Wetinator 2000) with small water reservoirs placed within.

## Single species experiments

### pH changing effect of collection of soil species

The collection of bacteria was isolated from a grain of local soil (Cambridge, MA, USA). The bacteria were pre-grown in TSB medium (Teknova, Hollister, USA) for 24h at RT in deepwell plates. The bacteria were diluted 1/100 into base medium with 10mM Phosphate and 8g/L urea and 10g/L glucose and grown for another 24 hours. The pH was measured in the wells that showed a final OD of >0.2 (100µL in 96well flat bottom plates). The same experiment was done by growing the bacteria in LB medium (Beckton Dickinson, Franklin Lakes, USA).

### pH changing effect of different species

Lp and Ca were precultured in 1xNu, pH7 and Sm and Pv were precultured in 1xNu, pH 7, 100mM PO_4_ overnight at 30°C. The next day Pv was diluted 1/100x and Sm 1/4000x into the same medium. Upon Pv and Sm reaching OD/cm 2 all four precultures were spun down with 4000rpm, 2min on a Eppendorf Tabletop centrifuge (Centrifuge 5810, with rotor A-4-81, Eppendorf, Hauppauge, USA) and washed two times with base buffer with 10mM phosphate pH7. The OD/cm was adjusted to 2. The bacteria were diluted 1/100x into 200µL base, pH 7, 10mM phosphate with 10g/L glucose and 8g/L urea in 96-deepwell plates (Eppendorf, Hauppauge, USA) and grown at 30°C and 1350rpm shaking speed for 24h. Afterwards the pH was measured in every well with the micro pH electrode (N6000BNC, SI Analytics, Weilheim, Germany).

### Growth of bacteria at different pH values

Lp and Ca were precultured in 1xNu, pH 7 and Sm and Pv were precultured in 1xNu, pH 7, 100mM PO_4_ overnight at 30°C. The next day Pv was diluted 1/100x and Sm 1/4000x into the same medium. Upon Pv and Sm reaching OD/cm 2 all four precultures were spun down with 4000rpm, 2min on the Eppendorf Tabletop centrifuge 5810 and washed two times with base buffer with 10mM Phosphate pH7. The bacteria were diluted 1/100x into 96-deepwell plates with 200µL of base, pH 7, 100mMPO4 in each well with a pH varying in integers from 2 to 11. The initial CFUs were estimated by plating the bacteria that should be incubated at pH 7 at the beginning of the incubation. After 24h at 30°C and 1350rpm shaking speed the bacteria were plated again and the final pH was measured (Supplementary Fig. 1C). The final CFU were divided by the initial CFU and the maximum was normalized to one to make the values comparable (Fig. 1B)

### Allee effect

The experiments were performed with *Corynebacterium ammoniagenes* (Ca) that is urease active and thus capable of cleaving urea into ammonia^19^. The bacterium was precultured in 2x5mL 1xNu with 100mM Phosphate, PH 7 in 50mL culture tubes at 250 rpm (Falcon/Becton Dickinson, Franklin Lakes, USA), 30°C for around 15 h. The bacteria were diluted 1/100x into 5mL of the same media and grown to an OD/cm of 2. The bacteria were spun down with 4000rpm on the Eppendorf tabletop centrifuge 5810 for 2 min and resuspended in base pH 6 with 10mM phosphate. The bacteria were spun down and resuspended again. The bacterial solution was split into two equal amounts, spun down and resuspended with 2.5mL of base pH 6 with either 10mM or 100mM phosphate. The OD/cm was adjusted to 2. For the experiments 96-well deepwell plates (Eppendorf, Hauppauge, USA) were used.

The experiment was performed in three different media: base, pH 6 with 10mM phosphate (low nutrient condition), base pH 6 with 10mM phosphate, 10g/L glucose and 8g/L urea (low buffer conditions) and base pH 6 with 100mM phosphate, 10g/L glucose and 8g/L urea (high buffer conditions). The final volume was 100µL per well. In the first row 270µL of media were prepared and 30µL of OD/cm = 2 bacteria solutions were added. Starting from this row 2/3 x (200µL into 100µL) dilutions were performed into the next rows. This way a gradient of initial cell densities was generated from the last row 200µL of bacteria solution were removed in the end to obtain 100µL in each well. The bacteria were cultivated at 30°C shaking at 1350rpm (Titramax 100, Heidolph North America, Elk Grove Village, USA). A 1/10x dilution into fresh medium was performed every 24h.

The CFU was estimated as described above immediately after setting up the experiments and after every 24 h growth cycle. Moreover the pH of the medium was measured at the end of the growth cycles. For every condition there were 4 replicates.

### Ecological suicide

For this experiment *Pseudomonas veronii* (Pv) was used, a bacterium that can alkalize the environment by urea cleavage but prefers itself lower pH for growth. The bacterium was grown at 30°C. The preculture of Pv was done in 5mL 1xNu, pH 7 with 100mM PO_4_ for around 15h. Pv was diluted 1/100x into the same medium and grown to an OD/cm of 2. The bacterial solution was washed two times with base with 10mM Phosphate, pH 7. The bacteria were resuspended in the same buffer and the OD/cm adjusted to 2. The 96-deepwell plates were prepared by adding 200µL base with 10mM Phosphate with or without 10g/L glucose and 8g/L urea (high and low nutrient conditions) or base with 100mM Phosphate with 10g/L glucose and 8g/L urea (high buffer condition) to the 96-well deepwell plates. The bacteria were added by 1/100x dilution. The plate was incubated at 30°C, 1350rpm shaking. Every 24h the bacteria were diluted 1/100x into fresh medium, the CFU estimated and the pH measured. For every condition there were 4 replicates.

## Interspecies interactions

### Bistability

*Corynebacterium ammoniagenes* (Ca) and *Lactobacillus plantarum* (Lp) were used here. Both species change the pH in a direction that they themselves like but the other species dislikes. Ca and Lp were pre-cultured for 17 h in 1xNu pH 7. The overnight cultures were washed with base with 10mM Phosphate pH 7 two times and the OD/cm adjusted to 2. The two bacterial species were mixed with ratios 1%, 10%, 30%, 45%, 55%, 70%, 90%, 99% keeping the sum OD/cm at 2. The bacterial mixture was diluted 1/100x into 96-deepwell plates containing 200µL base with 10mM Phosphate with/without 10g/L glucose and 8g/L urea. The starting pH was 7 for the medium without glucose and urea and 6, 6.5 and 7 for the medium with glucose and urea. The cultures were incubated at 30°C and shaken at 1350rpm. Every 24 h the CFU and pH were measured and the cultures were diluted 1/100x into fresh medium. For the CFU counting the dilutions of the cultures were plated onto TSB agar plates as described above. However, every dilution was plated twice once on a TSB agar plate with pH 5 and once with pH 10. At pH 5 only Lp could grow and at pH 10 only Ca, this way the bacteria could be easily distinguished. This was especially helpful since the colony morphology of Ca and Lp is rather similar (Supplementary Fig. 2). For every condition there were 3 replicates for the mixed culture and 8 replicates for the single species.

### Murder Suicide

*Lactobacillus plantarum* (Lp) and *Pseudomonas veronii* (Pv) were used for this experiment. Lp decreases the pH and likes low pH, whereas Pv increases the pH but prefers lower pH values. We made preculture of Lp in 1xNu pH 6 and Pv in 1xNu with 100mM phosphate pH 7 at 30°C 5ml in 50ml culture tubes. After around 15 h the bacteria were diluted 1/100x in the buffer they grew up in. Upon reaching OD/cm 2 the bacteria were spun down and washed two times with base with 10mM Phosphate, pH7. The OD/cm was adjusted to 2. The experiment was done in 96-deepwell plates each well containing 200µL base pH7 with 10mM or 100mM Phosphate and 10g/L glucose and 8g/L urea. The bacterial solution was diluted 1/100x (start OD/cm = 0.02) into the 96-deepwell plates. The plates were incubated at 30°C with 1350rpm shaking speed. Every 24h the CFU was estimated by plating, the pH was measured and the bacterial culture was diluted 1/100x into fresh medium. There were 8 replicates for every condition.

### Successive Growth

*Lactobacillus plantarum* (Lp) and *Serratia marcescens* (Sm) were used for this experiment. Sm was precultured in 1xNu with 100mM phosphate, pH 7, whereas Lp was precultured in 1xNutrient, pH6. After 15h preculture Sm was diluted 1/100x into the same buffer. Upon Sm reaching OD/cm 2 both cultures were spun down and washed two times with base with 10mM phosphate, pH7 and the OD/cm was adjusted to 2. Sm and Lp were mixed 1/1 resulting in a sum OD/cm of 2. The experiment was done in base with pH 10.2 and 10mM phosphate with and without 10g/L glucose or 100mM phosphate with 10g/L glucose. The glucose was added and the pH adjusted directly before setting up the experiments, followed by filtering the media. The bacteria were diluted 1/100x into the measurement medium (=0.02 OD/cm) and diluted 1/10x after every 24h cycle together with the measurement of the CFU and pH. There were 8 replicates for every condition.

### Mutual Stabilization

*Serratia marcescens* (Sm) and *Pseudomonas veronii* (Pv) were grown overnight in 1xNutrient with 100mM phosphate, pH 7 at 30°C. The next day Pv was diluted 1/100x and Sm 1/4000x into the same medium. Upon reaching OD/cm of 2 the bacteria were washed two times with base buffer with 10mM Phosphate pH 7. Afterwards the bacteria were resuspended in the same buffer and the OD/cm adjusted to 2. The bacteria were mixed 1/1 (vol/vol). The experiment was performed in 96-well deepwell plates with each well containing 200µL of base with 65mM phosphate, 10g/L glucose and 8g/L urea, pH 7. The bacterial mixture was diluted 1/100x into each well and the culture incubated at 30°C, shaken at 1350rpm. Every 24h the bacteria were diluted 1/10x into fresh medium and the bacteria were diluted 7 times 1/10x in PBS buffer. 150µL of the 10^-2^, 10^-4^, 10^-6^ dilutions were plated on TSB, 2.5% agar 100mm plates, each well on a full plate, for CFU measurement. Also the pH for every well was measured. There were 3 replicates for every condition.

## Reference

1. Atlas, R. M. & Bartha, R. Microbial ecology: Fundamentals and applications. (1986).

2. Faust, K. & Raes, J. Microbial interactions: from networks to models. Nat. Rev. Microbiol. 10, 538–550 (2012).

3. Fuhrman, J. A. Microbial community structure and its functional implications. Nature 459, 193– 199 (2009).

4. Raes, J. & Bork, P. Molecular eco-systems biology: towards an understanding of community function. Nat. Rev. Microbiol. 6, 693–699 (2008).

5. Strom, S. L. Microbial Ecology of Ocean Biogeochemistry: A Community Perspective. Science 320, 1043–1045 (2008).

6. Ghoul, M. & Mitri, S. The Ecology and Evolution of Microbial Competition. Trends Microbiol. 24, 833–845 (2016).

7. Shou, W., Ram, S. & Vilar, J. M. G. Synthetic cooperation in engineered yeast populations. Proc. Natl. Acad. Sci. 104, 1877–1882 (2007).

8. Nurmikko, V. Biochemical factors affecting symbiosis among bacteria. Experientia 12, 245–249 (1956).

9. Pande, S. et al. Fitness and stability of obligate cross-feeding interactions that emerge upon gene loss in bacteria. ISME J. 8, 953–962 (2014).

10. Mee, M. T., Collins, J. J., Church, G. M. & Wang, H. H. Syntrophic exchange in synthetic microbial communities. Proc. Natl. Acad. Sci. 111, E2149–E2156 (2014).

11. Hibbing, M. E., Fuqua, C., Parsek, M. R. & Peterson, S. B. Bacterial competition: surviving and thriving in the microbial jungle. Nat. Rev. Microbiol. 8, 15–25 (2010).

12. Yurtsev, E. A., Conwill, A. & Gore, J. Oscillatory dynamics in a bacterial cross-protection mutualism. Proc. Natl. Acad. Sci. 113, 6236–6241 (2016).

13. Bacteriocins: Evolution, Ecology, and Application. Annu. Rev. Microbiol. 56, 117–137 (2002).

14. Riley, M. A. & Gordon, D. M. The ecological role of bacteriocins in bacterial competition. Trends Microbiol. 7, 129–133 (1999).

15. Jones, R. T. et al. A comprehensive survey of soil acidobacterial diversity using pyrosequencing and clone library analyses. ISME J. 3, 442–453 (2009).

16. Rousk, J. et al. Soil bacterial and fungal communities across a pH gradient in an arable soil. ISME J. 4, 1340–1351 (2010).

17. Rousk, J., Brookes, P. C. & Bååth, E. Contrasting Soil pH Effects on Fungal and Bacterial Growth Suggest Functional Redundancy in Carbon Mineralization. Appl. Environ. Microbiol. 75, 1589– 1596 (2009).

18. Sofi, M. H. et al. pH of drinking water influences the composition of gut microbiome and type 1 diabetes incidence. Diabetes DB_130981 (2013).

19. Fu, W. & Mathews, A. P. Lactic acid production from lactose by Lactobacillus plantarum: kinetic model and effects of pH, substrate, and oxygen. Biochem. Eng. J. 3, 163–170 (1999).

20. Collins, M. Transfer of Brevibacterium ammoniagenes (Cooke and Keith) to the genus Corynebacterium as Corynebacterium ammoniagenes comb. nov. Int. J. Syst. Bacteriol. 37, 442– 443 (1987).

21. Solé, M., Rius, N. & Lorén, J. G. Rapid extracellular acidification induced by glucose metabolism in non-proliferating cells of Serratia marcescens. Int. Microbiol. 3, 39–43 (2010).

22. Allee, W. C. E. al. Principles of Animal Ecology. (W. B. Saunders, 1965).

23. Stephens, P. A., Sutherland, W. J. & Freckleton, R. P. What Is the Allee Effect? Oikos 87, 185–190 (1999).

24. Celiker, H. & Gore, J. Cellular cooperation: insights from microbes. Trends Cell Biol. 23, 9–15 (2013).

25. Gore, J., Youk, H. & van Oudenaarden, A. Snowdrift game dynamics and facultative cheating in yeast. Nature 459, 253–256 (2009).

26. Ratzke, C. & Gore, J. Self-organized patchiness facilitates survival in a cooperatively growing microbial population. Nature Microbiology, accepted (2016).

27. Drescher, K., Nadell, C. D., Stone, H. A., Wingreen, N. S. & Bassler, B. L. Solutions to the Public Goods Dilemma in Bacterial Biofilms. Curr. Biol. 24, 50–55 (2014).

28. Lahti, L., Salojärvi, J., Salonen, A., Scheffer, M. & Vos, W. M. de. Tipping elements in the human intestinal ecosystem. Nat. Commun. 5, 4344 (2014).

29. Datta, M. S., Sliwerska, E., Gore, J., Polz, M. F. & Cordero, O. X. Microbial interactions lead to rapid micro-scale successions on model marine particles. Nat. Commun. 7, 11965 (2016).

30. Koenig, J. E. et al. Succession of microbial consortia in the developing infant gut microbiome. Proc. Natl. Acad. Sci. 108, 4578–4585 (2011).

31. Klein, D. R. The Introduction, Increase, and Crash of Reindeer on St. Matthew Island. J. Wildl. Manag. 32, 350–367 (1968).

32. Hindell, M. A. Some Life-History Parameters of a Declining Population of Southern Elephant Seals, Mirounga leonina. J. Anim. Ecol. 60, 119–134 (1991).

33. Scheffer, V. B. The rise and fall of a reindeer herd. Sci. Mon. 73, 356–362 (1951).

34. Edwards, J. S., Ibarra, R. U. & Palsson, B. O. In silico predictions of Escherichia coli metabolic capabilities are consistent with experimental data. Nat. Biotechnol. 19, 125–130 (2001).

35. Kim, T. Y., Sohn, S. B., Kim, Y. B., Kim, W. J. & Lee, S. Y. Recent advances in reconstruction and applications of genome-scale metabolic models. Curr. Opin. Biotechnol. 23, 617–623 (2012).

36. Allen, E. E. & Banfield, J. F. Community genomics in microbial ecology and evolution. Nat. Rev. Microbiol. 3, 489–498 (2005).

37. Freilich, S. et al. Competitive and cooperative metabolic interactions in bacterial communities. Nat. Commun. 2, 589 (2011).

